# The Cystic Fibrosis treatment, Trikafta, affects growth, viability, and cell wall of *Aspergillus fumigatus* biofilms

**DOI:** 10.1101/2023.06.20.545692

**Authors:** Jane T. Jones, Kaesi A. Morelli, Elisa M. Vesely, Charles T.S. Puerner, Chetan K. Pavuluri, Brandon S. Ross, Norman van Rhijn, Michael J. Bromley, Robert A. Cramer

## Abstract

People with Cystic Fibrosis (pwCF) commonly test positive for the pathogenic fungus, *Aspergillus fumigatus,* which is associated with a decline in lung function. Trikafta is a recently approved therapy for pwCF that improves quantity and function of the CFTR protein, however, it is not known how Trikafta affects microbial communities in the lung. Therefore, the aim of this study was to determine whether Trikafta directly affects *A. fumigatus* growth and biology. While Trikafta did not impact the viability of *A. fumigatus* conidia, treatment of *A. fumigatus* biofilms with Trikafta reduced overall biofilm biomass. This finding was associated with increased membrane permeability, decreased viability, and reduced metabolic activity following long-term treatment of biofilms. Trikafta-induced membrane permeability and biomass reduction was partially blocked with the calcium channel inhibitor Verapamil and fully blocked by the mammalian CFTR inhibitor GlyH-101. Trikafta-induced biomass reduction and metabolic activity was shown to be regulated by the High Osmolarity Glycerol (HOG) pathway gene *sakA*. Trikafta treatment also induced resistance to the cell wall stressor calcofluor white, susceptibility to the antifungal caspofungin, and decreased inflammatory responses from murine bone marrow cells. Collectively, these results reveal that Trikafta affects infection-relevant *A. fumigatus* biology and host-microbial interactions.

**IMPORTANCE:** People with Cystic Fibrosis (pwCF) commonly test positive for the pathogenic fungus, *Aspergillus fumigatus*, which is associated with a decline in lung function. Trikafta is a recently approved therapy for pwCF that improves quantity and function of the CFTR protein, however, it is not known how Trikafta affects microbial communities in the CF lung. Here we report that Trikafta has anti-fungal activity against the infection relevant form of *A. fumigatus*, termed biofilms. When exposed to Trikafta *A. fumigatus* biofilms lose their viability. In addition, exposure to Trikafta enhances the susceptibility of *A. fumigatus* biofilms to the clinically utilized anti-fungal drug caspofungin. These data suggest Trikafta impacts the cell wall of A. fumigatus and suggests the potential for using Trikafta to augment existing anti-fungal therapy.

## INTRODUCTION

The triple drug combination therapy Trikafta (Elexacaftor/Tezacaftor/Ivacaftor) has been approved for Cystic Fibrosis patients harboring one or two copies of the dF508 *cftr* allele. The combination of two molecules involved in refining CFTR protein folding (Elexacaftor/Tezacaftor) and another molecule to potentiate CFTR ion gating function (Ivacaftor) has improved lung function, sweat chloride levels, and overall quality of life in pwCF (1, 2). Recent data has also shown that Elexacaftor can improve CFTR gating function in addition to folding (3). With the approval of Trikafta use in over 90% of the CF patient population, it is critical to understand the long-term impact of these combination treatments on the host and the associated microbial communities in the CF lung environment. Due to a thick buildup of sticky mucus, conventional host mechanisms that prevent microbial infection are perturbed in the CF lung and lead to chronic microbial colonization and infection. A wide variety of bacterial genera have been recovered from the CF lung, including *Pseudomonas, Streptococcus, Prevotella*, and many others. Additionally, diverse fungi (including *Candida spp*. and *Aspergillus spp*.) and viruses (influenza and respiratory syncytial viruses) have also been routinely identified (reviewed in: (4)).

The role of fungi in CF lung disease progression remains ill-defined. Due its frequent isolation from sputum samples of pwCF and its known role as a human fungal pathogen, the impact of *A. fumigatus* within the CF lung and subsequent pathogenesis is of particular importance. Invasive infections caused by *A. fumigatus* typically only occur in individuals with compromised immune systems, such as in chemotherapeutic or organ transplant patients (5–9). After *A. fumigatus* conidia germinate in the lung environment and form hyphae, a biofilm that is resistant to antifungal therapy develops (10, 11), which leads to high mortality (12–15). However, despite a functional immune system, *A. fumigatus* is detected in lung sputum samples of approximately 40-50% of pwCF and this colonization has been found to be associated with a sharper decline in lung function (16–18) compared to pwCF who test negative for *A. fumigatus*. Ivacaftor treatment is correlated with reduced *Aspergillus* species in the lungs of pwCF (19, 20) and a more recent publication showed that Orkambi (Ivacaftor/Lumacaftor) reduced phagocyte-induced ROS production from *A. fumigatus* (21). However, there have been no studies published directly addressing the potential impact of Trikafta on the biology of *A. fumigatus* itself and the potential consequences for the CF lung environment.

The overall goal of our study was to determine the impact of Trikafta on the biology of *A. fumigatus* and whether Trikafta treatment alters the host response to this important CF-associated fungus. We observed that Trikafta treatment has no minimum inhibitory concentration (MIC) against *A. fumigatus* conidia but does have a striking impact on biofilms in several laboratory and clinical isolates. Furthermore, our studies demonstrate that the effects of Trikafta on *A. fumigatus* biofilms are significantly exacerbated in combination with ionic stress. Collectively, we observed that Trikafta reduces the viability of *A. fumigatus* biofilms, increases membrane permeability, alters susceptibility to antifungal cell wall agents, and reduces the inflammatory response of murine bone marrow cells to *A. fumigatus*.

## RESULTS

*Trikafta reduces growth of both laboratory and clinical* A. fumigatus *biofilms*. To determine whether Trikafta directly affects *A. fumigatus* growth and/or function, we first tested whether Trikafta treatment affected conidial growth. Conidia from the two most utilized laboratory reference strains, CEA10 and Af293, were grown in either standard CLSI RPMI-1640 with MOPS buffer (22) or glucose minimal medium (GMM) with Trikafta doses ranging from 50μM/ individual modulator to 1.56μM/individual modulator. The antifungal voriconazole was used as a positive control. While the MIC value of both strains to voriconazole was 0.25 μg/ml, consistent with previously published data, Trikafta did not affect overall growth at any of the concentrations tested. We next tested whether Trikafta affected growth of *A. fumigatus* biofilms at different levels of maturity. Initiating (12hr), immature (16hr) and mature (24hr) CEA10 submerged biofilms were treated with either GMM plus DMSO (vehicle control; 0.15%) or Trikafta (5μM/molecule in every assay from this point forward) for an additional 24 hours before biomass was measured. Trikafta treatment of all stages of biofilm development reduced the total biomass, although only significantly in 12hr and 16hr biofilms (Fig 1A). Since biomass reduction was most pronounced at 16hr, 16hr biofilms were treated with either GMM plus DMSO control or Trikafta for 12hr and 24hr. Surprisingly, biomass reduction occurred in the Trikafta-treated biofilms compared to DMSO-treated controls only after 24hr treatment (Fig. 1B). To determine whether Trikafta effectively reduces biomass across a variety of *A. fumigatus* strains, we tested a selection of common laboratory strains (CEA10, Af293 and ATCC13073) as well as clinical isolates from Cystic Fibrosis respiratory samples (Af110-14.14, Af110-5.3, Af106-6.10). After 16hr of biofilm development, media was removed and biofilms were treated with GMM plus DMSO or Trikafta for an additional 24hr. Biofilms were imaged and dry biomass was quantified (Fig. 1C, D). For all strains tested, Trikafta significantly reduced total fungal biomass between 50-70%. Strikingly, the Trikafta treated biofilms macroscopically looked translucent and viscous, suggesting possible cell lysis had occurred (Fig. 1C).

**Figure 1:**
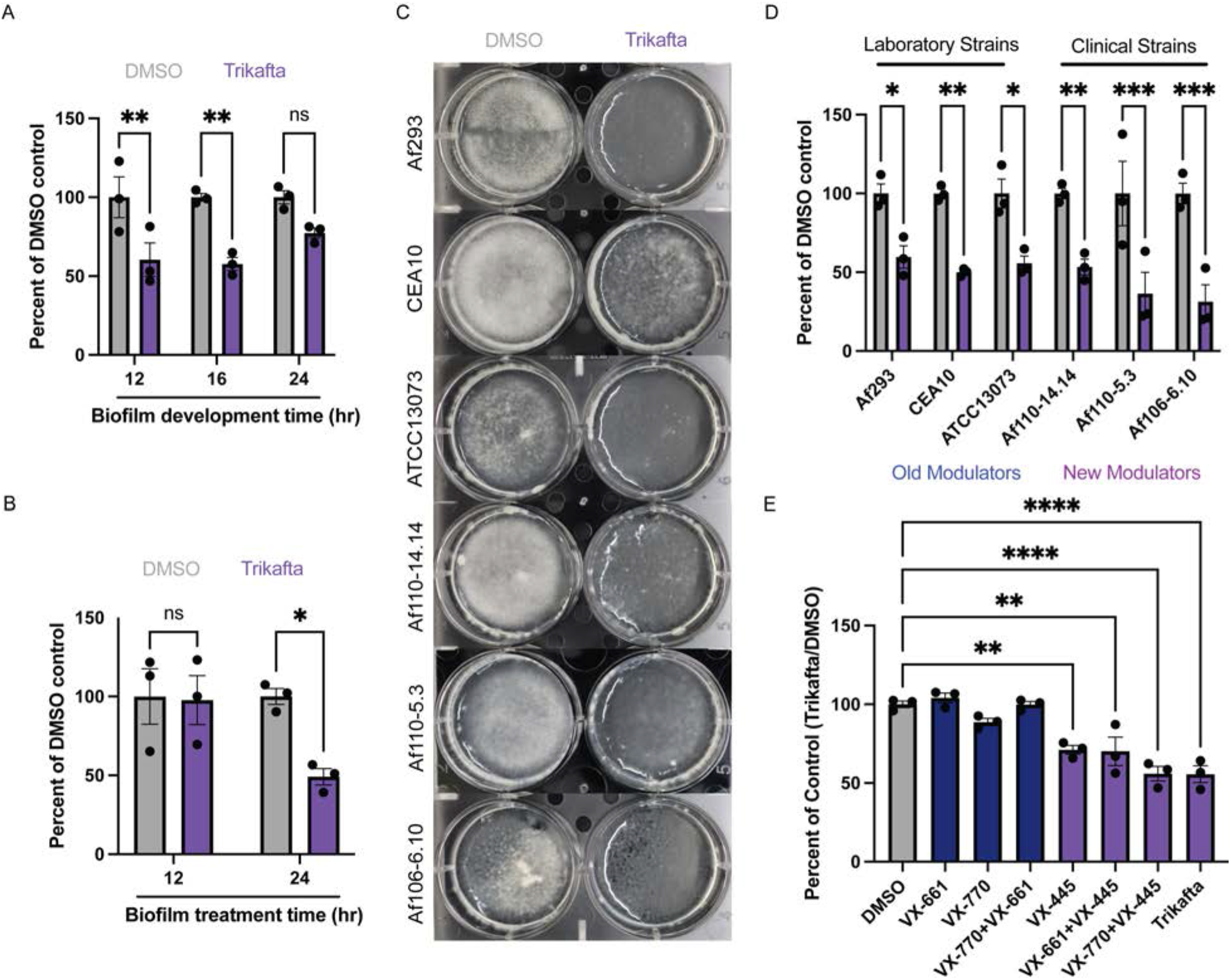
Trikafta reduces growth of both laboratory and clinical *A. fumigatus* biofilms. **(A)** CEA10 biofilms were grown in liquid glucose minimal media (L-GMM) for 12, 16 or 24hr. Media was removed and then fresh media with either DMSO (0.15%) or Trikafta (5μM/molecule) was added for an additional 24hr. Samples were collected, washed in water, and lyophilized. Dry biomass was measured, and all groups are represented as percent of DMSO-treated average. **(B)** CEA10 biofilms were grown in L-GMM for 16hr. Media was removed and then fresh media with either DMSO (0.15%) or Trikafta (5μM/molecule) was added for an additional 12 or 24hr. Samples were processed and graphically represented as in (A). **(C)** Biofilms of the indicated laboratory strains and clinical isolates were grown in L-GMM for 16hr. Media was removed and then fresh media with either DMSO (0.15%) or Trikafta (5μM/molecule) was added for an additional 24hr and images were taken. **(D)** Samples were processed and graphically represented as in (A). **(E)** CEA10 biofilms were grown in L-GMM for 16hr. Media was removed and then fresh media with either DMSO (0.15%), individual molecules, double combinations, or Trikafta (5μM/molecule) was added for an additional 24hr. Samples were processed and graphically represented as in (A). For (A, B, D), a 2-way ANOVA with Sidak’s multiple comparison was used and for (E) a one-way ANOVA with Dunnett’s multiple comparison with all groups compared to DMSO was used. Data points are three biological replicates, which represent the average of technical replicates.

Next we sought to define which molecule or combination of Ivacaftor (VX-770), Tezacaftor (VX-661), and Elexacaftor (VX-445) caused the reduction in fungal biomass. CEA10 16hr biofilms were treated with GMM plus DMSO, individual molecules, or different combinations of the individual molecules for an additional 24 hours before dry biomass was quantified (Fig. 1E). Single molecule Elexacaftor (VX-445) alone induced a significant reduction in total biomass (Fig. 1E). Interestingly, the Elexacaftor/Ivacaftor combination was as effective at reducing biomass as triple combination Trikafta, suggesting a potentially small but not significant effect for Ivacaftor as well. Collectively, these data show that Trikafta significantly affects growth of established biofilms, primarily through the action of the dual corrector/potentiator, Elexacaftor (VX-445).

*Trikafta reduces viability and increases membrane permeability of* A. fumigatus *biofilms*. Given the macroscopic observations of Trikafta-treated biofilms and the substantial loss of biomass, we sought to determine if Trikafta treatment reduced viability of the biofilms. To test this hypothesis, we determined whether Trikafta-treated biofilms could recover growth in fresh media. CEA10 biofilms were grown for 16hr and subsequently treated with GMM supplemented with either DMSO, Trikafta, or the antifungal drug Amphotericin B (AmB; 1μg/ml) as a positive control for an additional 1, 6, and 12hr. At each timepoint, baseline samples were collected and in parallel a recovery sample was prepared where media was removed and replaced with fresh GMM for a further 24hr of incubation before dry biomass weight was quantified (Fig. 2A). Biofilms treated with Trikafta and AmB for 1 and 6hr had around 50% of the recovery of DMSO treated biofilms. After 12hr treatment with Trikafta, biofilms had no recovery compared to three-fold recovery of DMSO-treated biofilms (Fig. 2B).

**Figure 2:**
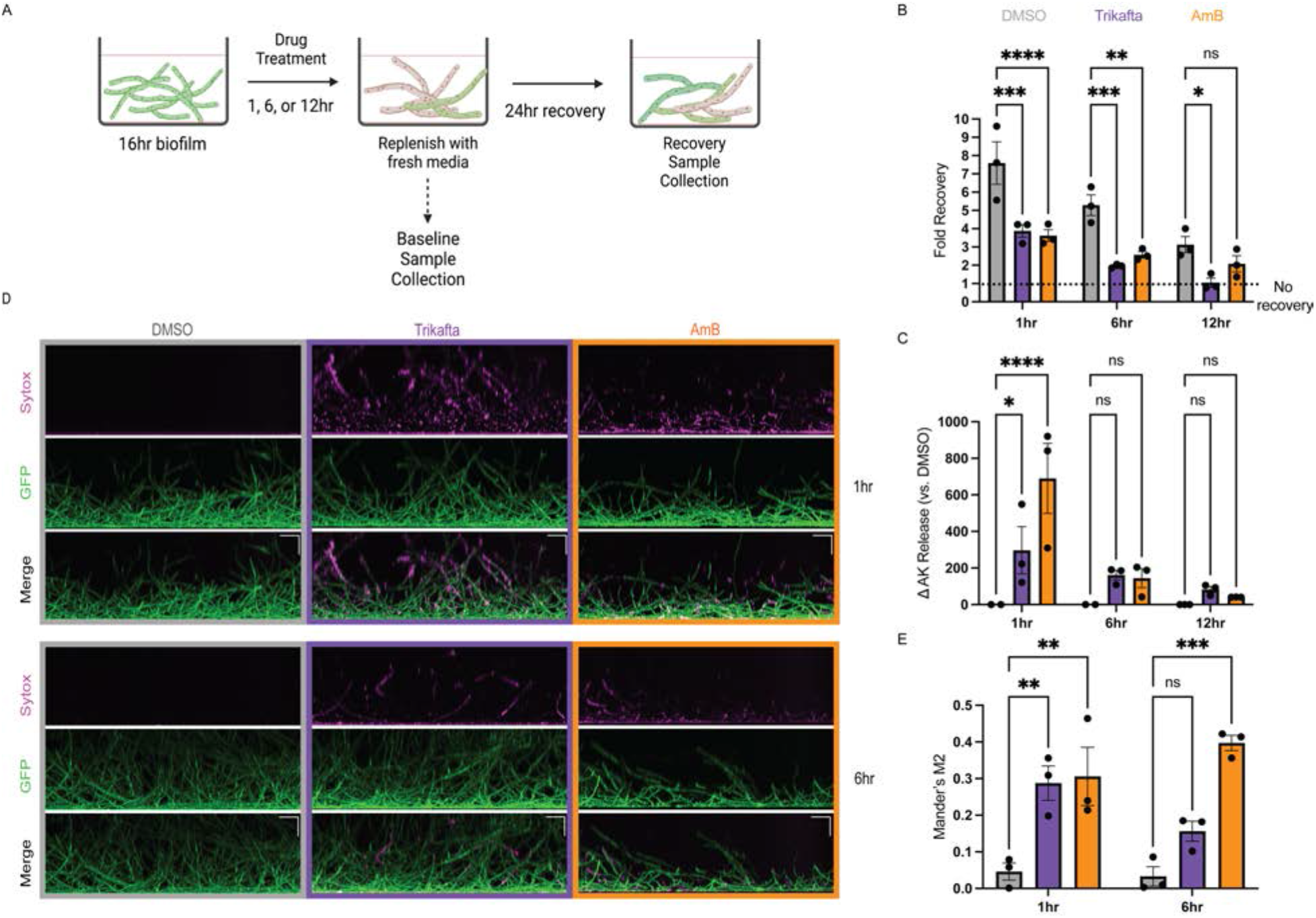
Trikafta reduces viability and increases membrane permeability of *A*. *fumigatus* biofilms. **(A)** Experimental design for viability assay **(B)** CEA10 biofilms were grown in L-GMM for 16hr. Media was removed and then fresh media with either DMSO (0.15%), Trikafta (5μM/molecule), or Amphotericin B (AmB) (1μg/ml) was added for an additional 1, 6, or 12hr. Baseline samples were collected, and parallel biofilms had media removed and then replaced with fresh L-GMM for an additional 24hr. Recovery biofilms were collected, washed in water, lyophilized, and dry biomass was measured. Treatment groups were compared to their own controls and data are represented as fold change from baseline control. **(C)** CEA10 biofilms were grown in L-GMM for 16hr. Media was removed and then fresh media with either DMSO (0.15%), Trikafta (5μM/molecule), or AmB (1μg/ml) was added for an additional 1hr, 6hr or 12hr. Supernatant was collected and adenylate kinase was measured. Data are shown as percent change over DMSO controls (DMSO set to 100%) and then subtracting 100 to get a delta in AK release. **(D)** CEA10-*gpdA:GFP* biofilms were grown in L-GMM for 16hr. Media was removed and then fresh media with either DMSO (0.15%), Trikafta (5μM/molecule), or AmB (1μg/ml) was added for an additional 1hr or 6hr. Sytox Blue 1:1000 was added for 5 minutes. Images were taken on Nikon spinning Disc confocal microscope 20X. **(E)** Images were quantified and represented as the fraction of GFP that overlaps with Sytox in each image (Mander’s M2). For (B, C, E), a 2-way ANOVA with Dunnett’s multiple comparison was used, with both treatment groups compared to DMSO controls. Data points are three biological replicates, which represent the average of technical replicates.

Next we determined whether Trikafta treatment affected overall membrane permeability of *A. fumigatus* biofilms. Supernatant from either DMSO or Trikafta-treated biofilms for 1, 6, or 12hr was analyzed for Adenylate Kinase (AK) activity, a marker of membrane permeability and damage (23). Treatment with Trikafta or AmB for 1hr significantly increased the change in AK release versus DMSO-treated controls approximately 300% and 600%, respectively. After 6 and 12hr treatment with Trikafta and AmB, detectable levels of AK were observed to around 50-100% change from DMSO controls, although to a much lesser extent than 1hr treatment (Fig. 2C). To further validate this observation, the cell viability stain, Sytox Blue, was used to image damage to biofilms. The stain was used in conjunction with a fluorescently labeled *A. fumigatus* strain constitutively expressing GFP in the cytoplasm. The GFP fluorescence allowed for identifying the overlap in Sytox signal. CEA10-*gpdA:GFP* conidia were grown into 16hr biofilms and treated with GMM plus either DMSO, Trikafta, or Amphotericin B for an additional 1 and 6hr prior to staining with Sytox Blue (12hr treatment was too long for interpretable microscopy images). Biofilms treated with GMM plus DMSO showed no intracellular Sytox Blue staining while biofilms treated with AmB showed around 30-40% positive staining at both 1 and 6hr treatment (Fig. 2D, E). Biofilms treated with Trikafta for 1hr were also around 30% positive for Sytox Blue and around 15% positive after 6hr of treatment (Fig. 2D, E). Collectively, these data demonstrate that Trikafta increases membrane permeability and reduces the viability of *A. fumigatus* biofilms.

*Trikafta modulates ion channels in* A. fumigatus *biofilms:* Trikafta modulates ion channel function in mammalian cells, so we next hypothesized that Trikafta-induced modulation of fungal ion channels causes membrane disruption and increased membrane permeability. We used a mammalian CFTR inhibitor GlyH-101 (24) and the calcium channel inhibitor Verapamil to address this hypothesis. CEA10-*gpdA:GFP* conidia were grown into 16h biofilms and treated with either DMSO control, GlyH-101 (20μM) or verapamil (1mM) for 1hr and then subsequently treated with Trikafta for an additional 1hr prior to Sytox Blue membrane permeability staining. Pre-treatment of biofilms with GlyH-101 and verapamil prior to Trikafta treatment almost completely abolished Trikafta-induced Sytox Blue staining (Fig. 3A, B). To test if this was specific to Trikafta, we performed the same set of experiments with Amphotericin B. Pre-treatment with GlyH-101 and verapamil had little to no effect on Amphotericin B-induced Sytox staining (Fig. 3A, B). To determine whether ion channel modulation contributes to Trikafta-induced reduction in biomass, 16hr biofilms were pre-treated with GlyH-101 or verapamil for 1hr then treated with Trikafta or Amphotericin B for 24 hr. GlyH-101 almost fully rescued the biomass reduction of Trikafta while verapamil partially rescued it (Fig. 3C). GlyH-101 and Verapamil had no effect on Amphotericin B-induced biomass reduction (Fig. 3C). Therefore, the impact of ion channel inhibitors on Trikafta are likely unique to its mechanism of action, not just general membrane stress since the inhibitors did not affect AmB-induced effects.

**Figure 3:**
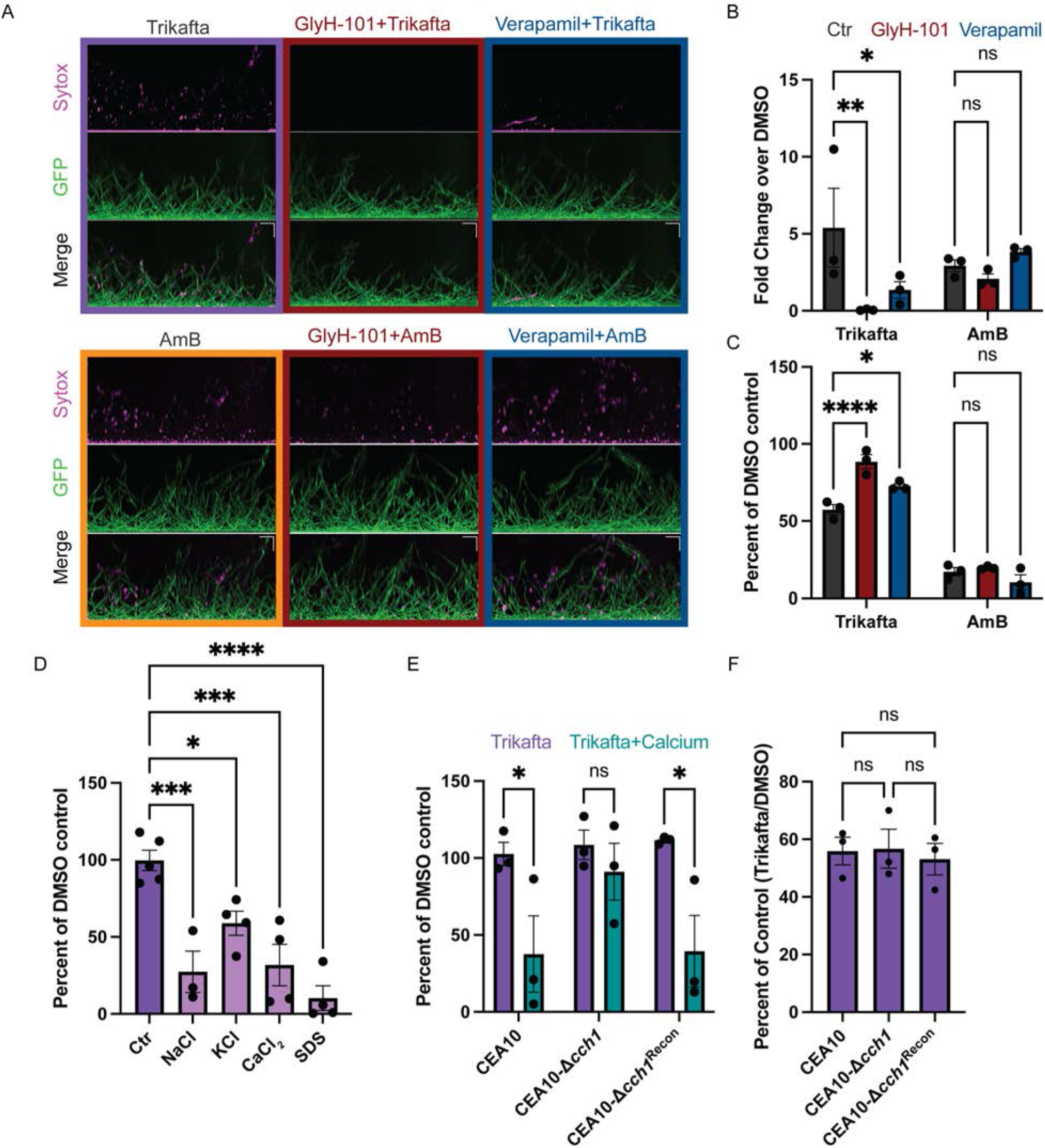
Trikafta modulates ion channels in *A. fumigatus* biofilms. **(A)** CEA10-*gpdA:GFP* biofilms were grown in L-GMM for 16hr. Media was removed and biofilms were treated with DMSO (control, 0.2%), Verapamil (calcium channel inhibitor, 1mM), or GlyH-101 (CFTR inhibitor, 20μM) for 1hr. Media with either DMSO (0.15%), Trikafta (final concentration 5μM/molecule) or AmB (final concentration 1μg/ml) was added for an additional 1hr. Sytox Blue 1:1000 was added for 5 minutes. Images were taken on Nikon spinning Disc confocal microscope at 20X. **(B)** Images were quantified and represented as the fraction of GFP that overlaps with Sytox in each image (Mander’s M2) and data are represented as fold change over DMSO-treated control. **(C)** CEA10 biofilms were grown for 16hr and media removed. Fresh media with either DMSO (0.2%), GlyH-101 (20μM) or Verapamil (1mM) was added for 1hr. Subsequently, Media with either DMSO (0.15%), AmB (Final dose 1μg/ml), or Trikafta (Final dose 5μM/molecule) was added for an additional 24h. Samples were processed and graphically represented as in Fig 1A. **(D)** CEA10 biofilms were grown in L-GMM for 16hr. Media was removed and then fresh media with either DMSO (0.15%) or Trikafta (5μM/molecule) was added in either vehicle control (H_2_0) or 0.5M NaCl, CaCl_2_, or KCl or 0.002% SDS for an additional hour. Metabolic activity was measured using XTT assay. Data are normalized as percent of Trikafta/DMSO controls in each condition. **(E)** WT CEA10, CEA10-Δ*cch1*, and CEA10-*cch1^rec^* biofilms were grown in L-GMM for 16hr. Media was removed and then fresh media with either DMSO (0.15%) or Trikafta (5μM/molecule) was added in either control or 0.5M CaCl_2_for an additional hour and an XTT assay was performed. Data are normalized as percent of Trikafta/DMSO controls in either vehicle control (H_2_O) or CaCl_2_. **(F)** WT CEA10, CEA10-Δ*cch1*, and CEA10-*cch1^rec^* biofilms were grown in L-GMM for 16hr. Media was removed and then fresh media with either DMSO (0.15%) or Trikafta (5μM/molecule) was added for an additional 24hr. Samples were processed and graphically represented as in Fig 1A. For B, C, and E, a 2-way ANOVA with Dunnett’s (B and C) or Sidak’s (E) multiple comparisons was performed and a one-way ANOVA (D and F) with Dunnett’s (D) or Tukey’s (F) multiple comparisons was performed. Data points are three biological replicates, which represent the average of technical replicates.

We next determined whether the effects of Trikafta were exacerbated in the presence of ionic stress. We chose to utilize an XTT-based metabolic assay to measure the metabolic activity of Trikafta-treated biofilms after 1hr treatment because there was little to no effect of Trikafta on metabolic activity at that time point (Fig. 3D). Therefore, we treated 16hr biofilms with either GMM plus DMSO control or Trikafta for 1hr in either control, 0.5M NaCl, 0.5M CaCl_2_, or 0.5M KCl. In parallel, we also included 0.002% SDS as a control for general membrane disruption. It is important to note that the low dose of SDS used did not cause any reduction in metabolic activity on its own in contrast to the different ions (data not shown). Strikingly, Trikafta treatment reduced metabolic activity by more than 70% in the presence of NaCl, CaCl_2_, and SDS. Trikafta also reduced metabolic activity by more than 40% in the presence of KCl (Fig. 3D). We next hypothesized that a strain harboring a deletion mutation in the calcium channel gene, *cch1* (25), would be resistant to Trikafta-reduced metabolic activity in CaCl_2_ stress conditions. CEA10-ι1*cch1* biofilms treated with Trikafta in the presence of CaCl_2_ were resistant to reduced metabolic activity compared to WT CEA10 biofilms (Fig 3E). However, the CEA10-Δ*cch1* biofilms treated with Trikafta for 24hr had roughly equal susceptibility to Trikafta as WT controls, indicating that exogenous CaCl_2_ availability may be required (Fig 3E). These data suggest that the effects of Trikafta are exacerbated in ionic stress conditions.

*Trikafta transiently increases metabolic activity in A. fumigatus biofilms*: Since the CEA10-Δ*cch1* mutant biofilm was not resistant to Trikafta-induced biomass reduction, we further sought to determine how Trikafta reduced biofilm biomass after long-term exposure to the drug. To establish an assay for further exploring the metabolic changes in fungal biomass, we evaluated biofilms over time using the metabolic dye Resazurin, as it allows for data collection over a range of time rather than a snapshot in time as with XTT-based metabolic assays. We treated 16h biofilms with either GMM plus DMSO or Trikafta for 30 minutes and then added Resazurin for an additional 5hr. Surprisingly, analysis of metabolic activity in Trikafta-treated biofilms compared to DMSO-treated controls showed a 30-40% increase in metabolic activity in CEA10 biofilms (Fig. 4A). We hypothesized that this increased metabolic activity was likely due to increased mitochondrial activity.

**Figure 4:**
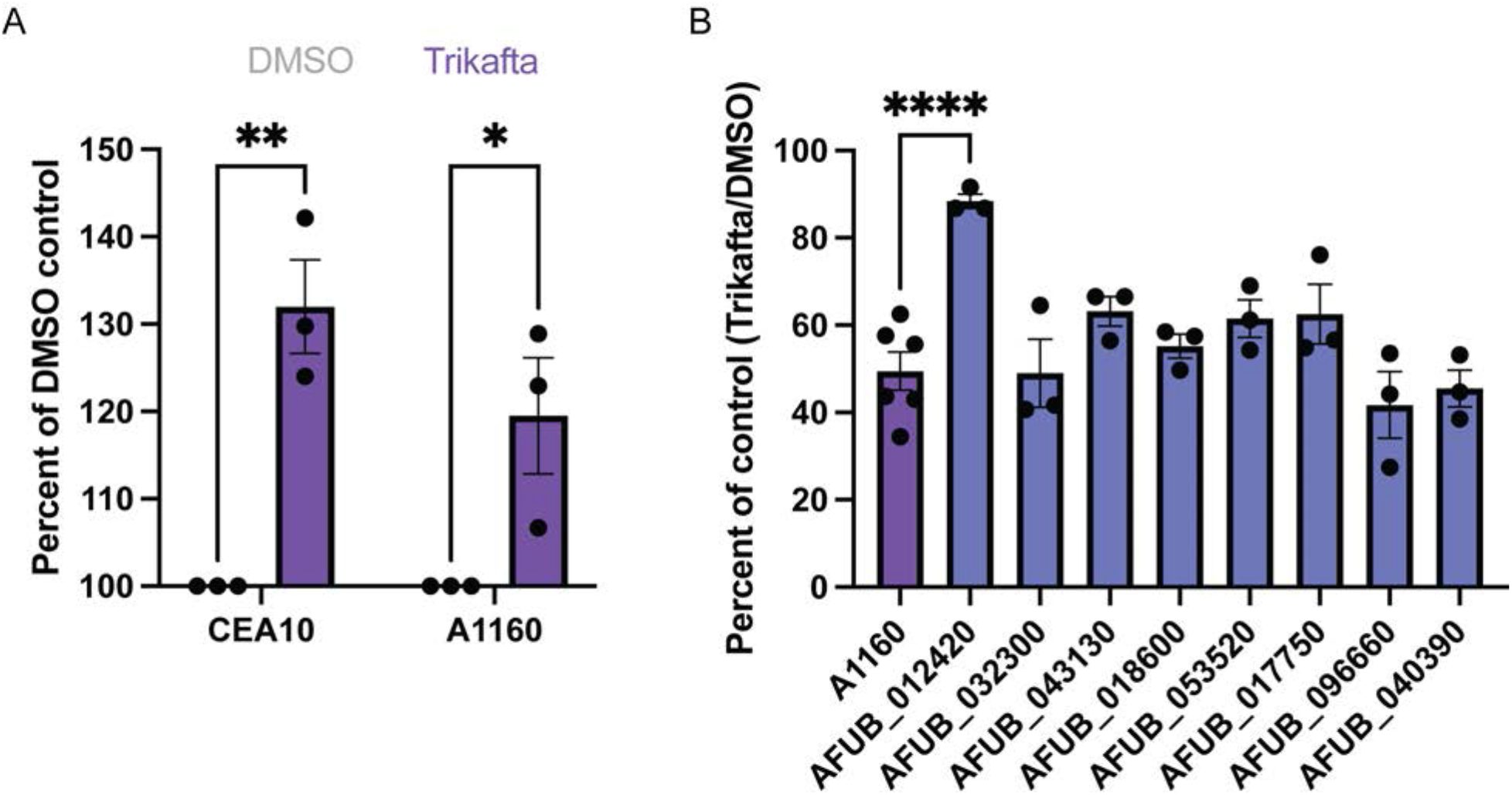
Trikafta acutely increases metabolic activity in *A. fumigatus* biofilms. **(A)** CEA10 or A1160 biofilms were grown in L-GMM for 16hr. Media was removed and then fresh media with either DMSO (0.15%) or Trikafta (5μM/molecule) was added for an additional 30 minutes. The metabolic dye, Resazurin, was added 10% volume for an additional 5hr after Trikafta treatment and fluorescence 594 was captured. A 2way ANOVA with Sidak’s multiple comparisons was used. Data points are three biological replicates, which represent the average of technical replicates. **(B)** Biofilms of the indicated kinase library mutant strains were grown in L-GMM for 16hr. Media was removed and then fresh media with either DMSO (0.15%) or Trikafta (5μM/molecule) was added for an additional 24hr. Samples were processed and graphically represented as in Fig 1A. 3-6 technical replicates are represented.

To begin to delineate pathway(s) by which Trikafta acts on *A. fumigatus* biofilm biomass reduction, we used this Resazurin assay to perform a genetic screen on an *A. fumigatus* kinase knock-out library. Importantly, we observed a similar increase in metabolic activity in the WT A1160 background strain used to generate the knockout library upon Trikafta treatment (Fig 4A). A total of 108 putative kinases were grown into 16hr biofilms, treated with either GMM plus DMSO or Trikafta for 30 minutes and subsequently treated with Resazurin for an additional 5hr. For each run, the effect of Trikafta-induced metabolic activity in the parent strain (A1160) was calculated and used as the reference. A total of 18 strains had altered metabolic activity in response to Trikafta compared to the control strain, based upon being either 20% below or 15% above the control strain (Table 2). Eight mutant strains with the highest change in susceptibility were selected to screen for the biomass phenotype and it was found that a strain with a deletion mutation in AFUB_012420 (high osmolarity glycerol (HOG) pathway kinase, *sakA*) was the least susceptible to Trikafta-induced biomass reduction (Fig. 4B).

*Trikafta activates SakA and reduction of SakA activity increases resistance of A. fumigatus biofilms to Trikafta.* To confirm if loss of *sakA* alters biofilm biomass reduction to Trikafta, we next tested A1160-WT, A1160-1′*sakA*, Afs35-WT and Afs35-1′*sakA* biofilms that were grown for 16hr and treated with either DMSO or Trikafta for an additional 24 hours prior to imaging (Fig. 5A). Quantification of biomass showed that while both WT (A1160 and Afs35) biofilms had approximately 50-60% reduction in biomass with Trikafta treatment compared to controls, both Δ*sakA* mutant biofilms had little to no reduction in biomass with Trikafta treatment (Fig. 5B). We also observed that A1160-1′*sakA* biofilms were resistant to Trikafta-induced AK release after 12hr treatment (Fig. 5C). To further determine the effects of Trikafta at this later time point, 16hr A1160 and A1160-1′*sakA* biofilms were treated with either DMSO or Trikafta for 12hr and metabolic activity was assessed by XTT reduction. Excitingly, Trikafta caused reduction of metabolic activity by 50% in WT A1160, while A1160-1′*sakA* biofilms only had an approximately 20% reduction in metabolic activity (Fig. 5D). This further supports our hypothesis that the initial metabolic burst observed in response to Trikafta is followed by a marked reduction in metabolic activity and that Trikafta-induced changes in metabolic activity are regulated by *sakA*.

**Figure 5:**
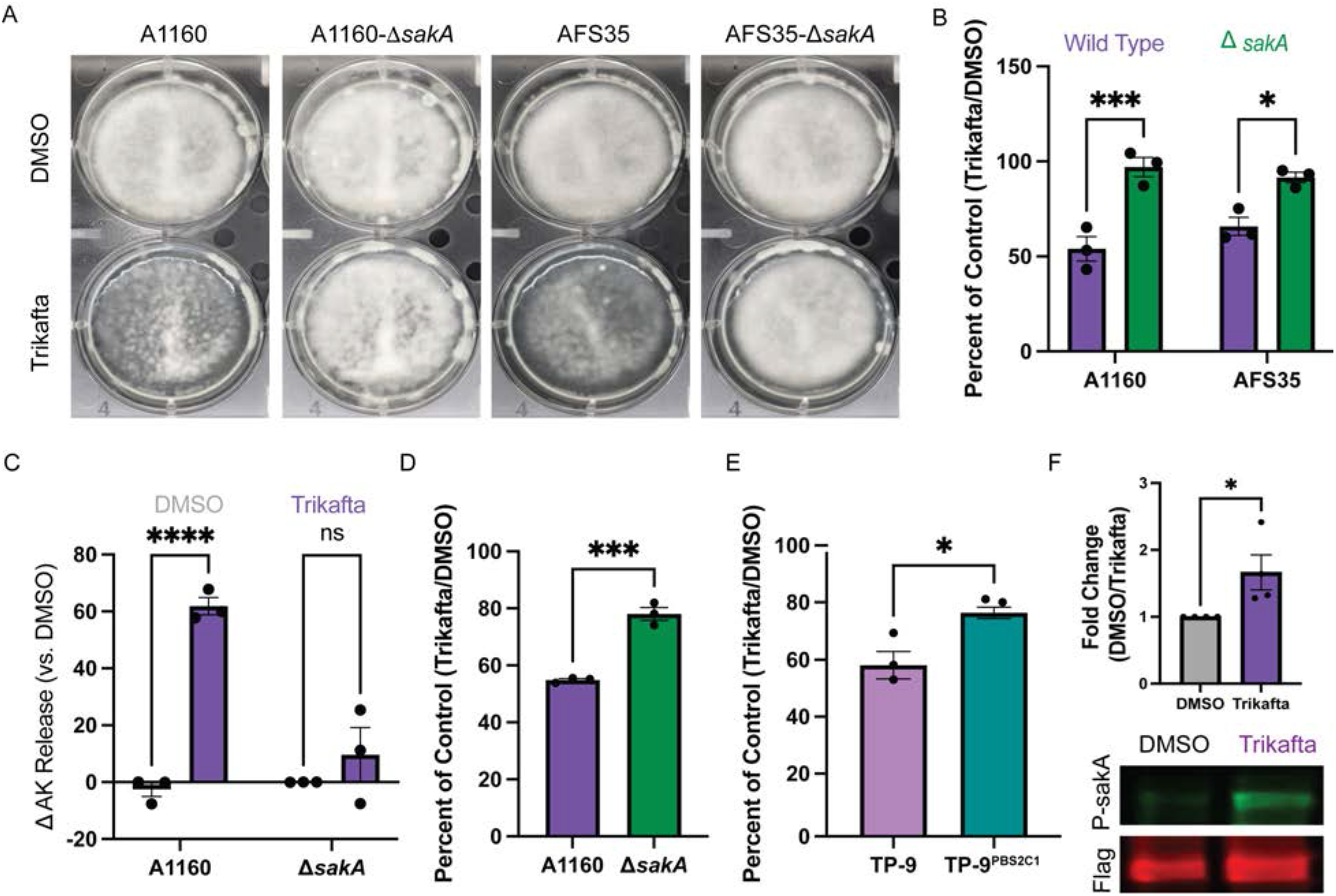
Trikafta activates sakA and reduction of sakA activity increases resistance of *A. fumigatus* biofilms to Trikafta. **(A)** WT (A1160 and Afs35) and Δ*sakA* conidia (in both WT strains) were grown for 16hr into biofilms. Media was removed and then fresh media with either DMSO (0.15%) or Trikafta (5μM/molecule) was added for an additional 24hr and images were taken. **(B)** Samples were processed and graphically represented as in Fig 1A. **(C)** WT A1160 and A1160-Δ*sakA* biofilms were grown in L-GMM for 16hr. Media was removed and then fresh media with either DMSO (0.15%) or Trikafta (5μM/molecule) was added for an additional 12hr. Supernatant was collected and Adenylate Kinase was quantified. Data are shown as percent change over DMSO controls (DMSO set to 100%) and then subtracting 100 to get a delta in AK release. **(D)** In parallel to (C), metabolic activity of biofilms was measured by XTT. Data are represented as percent of DMSO controls (Trikafta/DMSO). **(E)** TP9 and TP-9^PBS2C1^ biofilms were grown for 16hr. Media was removed and then fresh media with either DMSO (0.15%) or Trikafta (5μM/molecule) was added for an additional 24hr. Samples were processed and graphically represented as in Fig 1A. **(F)** TP-9^PBS2C1: sakA^ ^Flag^ biofilms were grown in L-GMM for 16hr. Media was removed and then fresh media with either DMSO (0.15%) or Trikafta (5μM/molecule) was added for an additional 30 minutes. Biofilms were collected and protein extracted for western blot analysis of either total sakA (Flag) or P-sakA. Samples were normalized to total protein and quantified using Licor. A 2-way ANOVA with Sidak’s multiple comparison (B and C) and a student’s t-test (D-F) were performed. Data points are three biological replicates, which represent the average of technical replicates.

Previous work from our laboratory has shown that an *A. fumigatus* clinical isolate from a pwCF, designated TP-9, harbors mutations in the MAP kinase kinase *pbs2* that regulates SakA pathway activity. This unique *pbs2* allele arose from long term growth in the CF lung environment and functionally leads to increased activation of SakA as indicated by phosphorylation (26). An allele swap with a *pbs2* allele from a different *A. fumigatus* clade isolate from the same patient generated the strain TP-9^PBS2C1^ and results in reduced SakA activation compared to TP-9 (26). To further test the effects of SakA activity on the response to Trikafta, we next compared Trikafta-induced effects on these two strains. TP-9 and TP-9^PBS2C1^ biofilms were grown for 16hr and treated with GMM plus either DMSO or Trikafta for an additional 24hr. Quantification of biofilm biomass showed that while TP-9 biofilms had almost 45% reduction in biomass with Trikafta treatment compared to controls, TP-9^PBS2C1^ only had 20% reduction, further showing that a reduction in SakA activity conferred resistance to Trikafta-induced biomass reduction (Fig. 5E). Additionally, we wanted to confirm if Trikafta induced phosphorylation of SakA. Utilizing the TP-9^PBS2C1:sakA-Flag^ strain (26), 16hr biofilms were treated with GMM plus either DMSO or Trikafta for 30 minutes, biomass was collected, and protein subjected to SDS-PAGE and subsequent Western Blot analysis for total SakA (Flag) and P-SakA. Trikafta treatment increased P-SakA almost two-fold in comparison to DMSO-treated controls, suggesting that Trikafta treatment activates SakA in *A. fumigatus* biofilms (Fig 5F).

*Trikafta modulates the biofilm cell wall and mediates inflammatory responses in host immune cells.* Since Trikafta treatment takes place in the complex CF lung environment, we next explored how Trikafta co-treatment with stress agents and conditions found in the CF lung environment affects *A. fumigatus* biofilms. We co-treated 16hr CEA10 biofilms with either DMSO or Trikafta with increasing doses of antifungal drugs, cell wall stress agents, metabolic stress agents, and oxidative stress inducers and measured overall metabolic activity as a readout of fungal cell damage using the XTT assay (27) (Fig. 6A, B). Trikafta alone did not affect metabolic activity of biofilms compared to DMSO controls after 3 hours of treatment (data not shown). Voriconazole, an antifungal drug targeting ergosterol biosynthesis, only mildly damaged the biofilms by about 20-30%, as previously published (12). Intriguingly, Trikafta modestly protected biofilms from voriconazole-induced damage at this time point. Trikafta co-treatment with Amphotericin B, which damages the cell membrane through direct binding of ergosterol, increased fungal cell damage. The effects of the echinocandin β-(1,3)-glucan synthase inhibitor, caspofungin, were exacerbated by Trikafta at a lower dose (0.015µg/ml). Trikafta also exacerbated the effects of mitochondrial inhibitors FCCP and antimycin A. Unexpectedly, Trikafta strongly protected biofilms from calcofluor white (CFW)-induced damage (Fig. 6A, B). Since Trikafta had strong impacts on cell wall stressors CFW and caspofungin, we decided to confirm our screen results. We observed that Trikafta protected biofilms from CFW-induced damage (Fig. 6C) and exacerbated a low dose of caspofungin-induced damage (Fig. 6D), confirming Trikafta altered biofilm responses to cell wall stressors.

**Figure 6:**
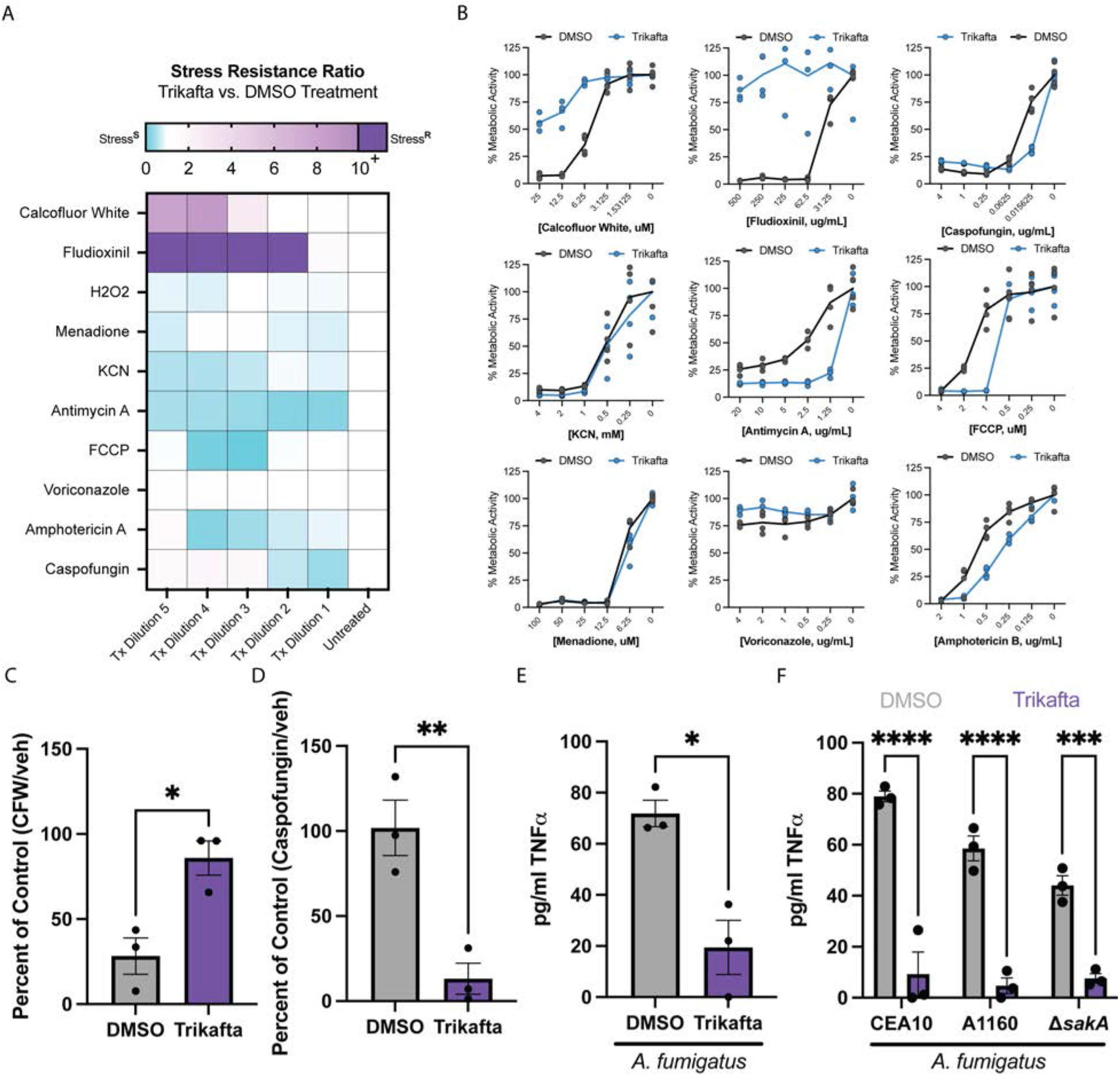
Trikafta modulates the biofilm cell wall and mediates inflammatory responses in host immune cells. **(A)** CEA10 biofilms were grown in L-GMM for 16hr. Media was removed and then fresh media with either DMSO (0.15%) or Trikafta (5μM/molecule) plus serial dilutions of the indicated stressors (CFW, Fludioxinil, caspofungin, KCN, Antimycin A, FCCP, menadione, voriconazole, and AmB) was added for an additional 3 hrs. Metabolic activity was measured by the reduction of the dye, XTT. Stress resistance ratio (Trikafta/DMSO controls) is represented by highly resistant (purple) through highly susceptible (blue) as a consequence of Trikafta co-treatment. **(B)** Graphs represent the percent of stressor/control in the presence or absence of Trikafta. Data represent 5 technical replicates. **(C)** CEA10 biofilms were grown in L-GMM for 16hr. Media was removed and then fresh media with either DMSO (0.15%) or Trikafta (5μM/molecule) plus 12.5μM CFW or **(D)** 0.015625μg/ml caspofungin for three hours. An XTT assay was performed to determine metabolic activity and data are shown as percent of stressor/control in either DMSO or Trikafta. **(E)** Bone marrow cells were isolated from C57Bl/6 mice harboring dF508 mutation in *cftr* and seeded 1X10^7/ml in co-culture with CEA10 conidia (1X10^6/ml) for 16hr in the presence of IFNγ (10ng/ml) with either DMSO (0.15%) or Trikafta (5μM). 96 well plates were centrifuged and supernatants were analyzed for murine TNFα. **(F)** Bone marrow cells were isolated from WT C57Bl/6 mice and seeded 1X10^7/ml in co-culture with CEA10, A1160, or A1160-Δ*sakA* conidia (1X10^6/ml) for 16hr in the presence of IFNγ (10ng/ml) with either DMSO (0.15%) or Trikafta (5μM). 96 well plates were centrifuged and supernatants were analyzed for murine TNFα. A student’s t-test was performed for C-E and a 2way ANOVA with Sidak’s multiple comparisons was performed for F. For C-F data points are three biological replicates, which represent the average of technical replicates.

Since Trikafta protected biofilms against CFW but exacerbated caspofungin effects, we hypothesized that Trikafta was likely increasing chitin and decreasing (1,3)-β-glucan content of the fungal cell wall. Furthermore, it has previously been shown that altering chitin content results in a compensatory effect that increases levels of (1,3)-ý-glucan (28). Since mammalian cells recognize (1,3)-β-glucan, which triggers an inflammatory response, we hypothesized that Trikafta treatment would reduce the inflammatory response of host cells in co-culture with *A. fumigatus*. To test this hypothesis, CEA10 conidia were co-cultured with bone marrow (BM) cells from mice harboring the mouse dF508 mutation in both *cftr* alleles (29) for 16hr. Supernatant was removed and assayed for TNFα concentrations. Co-culture of murine bone marrow cells with *A. fumigatus* elicited an increase in TNFα production, however, this effect was significantly reduced by Trikafta treatment (Fig 6E). Finally, we tested whether Trikafta altered inflammatory cytokine production from WT BM cells. Excitingly, Trikafta treatment also reduced TNFα production from WT BM cells in co-culture with *A. fumigatus* compared to vehicle controls (Fig. 6F). Furthermore, A1160 and A1160-1′*sakA* were equally susceptible to Trikafta-induced protection against the inflammatory response elicited in co-culture with murine BM cells, suggesting that the effects of Trikafta on the cell wall are upstream of SakA-mediated biofilm biomass reduction (Fig. 6F). These data collectively demonstrate that the effects of Trikafta alter the fungal cell wall, which subsequently change how host cells respond to *A. fumigatus,* independent of *cftr* mutations.

## DISCUSSION

The objective of this study was to determine if the current state-of-the-art treatment for pwCF, Trikafta, affects the biology of a common CF associated fungus, *A. fumigatus*. To begin to address this question, standard microbroth dilution MIC assays were utilized, and no antifungal effect was observed. However, the MIC assay tests the effects of a drug against conidia, so we tested whether Trikafta affected *A. fumigatus* biofilms, which are found in an established infection. Unexpectedly, we observed that long term (24hr) treatment of biofilms with Trikafta reduced *A. fumigatus* biomass in several laboratory and clinical strains (Fig. 1C, D). The reduction in biofilm biomass correlated with reduced cell viability and increased membrane permeability (Fig 2). Interestingly, the effects of Trikafta on permeability and viability occurred acutely, whereas biofilm biomass decrease was not observed until between 12-24hr post treatment (Fig. 1B).

Surprisingly, there are only three major classes of approved antifungal drug therapies for the treatment of aspergillosis: polyenes, echinocandins, and azoles (30). Polyenes and azoles target the fungal cell membrane and echinocandins target the fungal cell wall. Our data strongly suggest that Trikafta reduces viability in *A. fumigatus* biofilms by increasing membrane permeability and subsequent metabolic dysfunction. Within 1hr of treatment with Trikafta, biofilms showed a significant increase in Sytox Blue staining and AK release (Fig 2) and exacerbation of SDS-induced cell damage (Fig. 3D). These effects occur in parallel to Trikafta’s impact on overall viability of the biofilm. AK, which is highly expressed in the cytosol, is released extracellularly after the fungal cell integrity is compromised. It is also secreted when *A. fumigatus* is not actively growing or germinating (23). AK release has been successfully leveraged to identify antifungal compounds in several fungal species, such as *A. fumigatus* (23), *Candida albicans* (31), and *Cryptococcus neoformans* (32). Although Trikafta does not reduce overall growth of biofilms to the same magnitude as AmB treatment (Fig 3C), we observed a striking effect of Trikafta on biofilm membrane permeability, viability, and growth. Considering the drug resistant nature of fungal biofilms, future studies examining how these observations are relevant in the context of *in vivo*-relevant stressors are warranted.

Our data strongly suggest that Trikafta modulates ion channels in *A. fumigatus* biofilms. Given that treatment with Trikafta took more than 12hr to affect biofilm growth, we were surprised to observe membrane permeabilization and reduced viability within 1hr of treatment (Fig. 2A-C). Since we observed an effect on the cell membrane, we hypothesized that Trikafta’s mode of action is potentially like its function in mammalian cells. Previous studies in the literature have shown that calcium-activated chloride channels can also act as dual scramblases, which alter membrane fluidity and function (33, 34). Verapamil has been successfully utilized to block calcium channels in *Aspergillus fumigatus* (35) and our data demonstrate that it inhibits Trikafta-induced membrane permeability and biomass reduction (Fig. 3A and 3B). Our data also support the hypothesis that Trikafta may target multiple ion channels, due to its exacerbation in conditions with high NaCl, CaCl_2_, and KCl levels (Fig. 3C). Since our null mutant for *cch1* is resistant to Trikafta damage in calcium stress, this lends support that there are multiple ion channel targets for *A. fumigatus*, particularly since the CEA10-Δ*cch1* strain is equally susceptible to Trikafta-induced biomass reduction as WT (Fig. 3E). We also searched for CFTR-like proteins using the human *cftr* sequence in the *A. fumigatus* genome and identified eight highly related sequences, with the top hit being AFUB_066250. This gene is a putative ABC multidrug transporter with orthologs shown to have roles in secondary metabolite biosynthetic processes. We generated a null mutant in this gene and did not observe differences in Trikafta-induced biomass reduction compared to WT and reconstituted controls (data not shown).

In addition to effects on membrane permeabilization and ion channel modulation, our data also suggest that Trikafta affects cell wall composition in fungal biofilms. Interestingly, when biofilms were exposed to Trikafta and lower concentrations of caspofungin, Trikafta increased biofilm susceptibility to caspofungin-induced damage (Fig. 6A, B, and D). Alternatively, Trikafta protected biofilms from high doses of caspofungin (Fig. 6A, B). When biofilms were exposed to Trikafta and the cell wall stressor, CFW, Trikafta protected the biofilms from CFW-induced damage (Fig. 6A-C). Since CFW directly binds and blocks chitin polymerization and higher doses of caspofungin induce chitin biosynthesis via the paradoxical effect (36), it is plausible that Trikafta increases chitin composition in the biofilm cell wall. The caspofungin paradoxical effect (CPE) occurs when (1,3)-β-glucan synthase is inhibited to such a degree that the cell compensates by increasing chitin content. CPE-induced chitin synthesis has been shown to be stimulated via the cell wall integrity pathway mitogen-activated protein kinase (MAPK) signaling cascade and the transcription factors RlmA and CrzA in *A. fumigatus* strain Af293 (37). However, this effect has been shown to be strain-specific, as CrzA is not required for CPE in CEA10-derived strains (37).

Furthermore, depletion of (1,3)-β-glucan synthases results in increased chitin in *Candida albicans*, which is dependent on calcium (38). Future studies examining the effects of calcium, CPE, and CWI pathway are critical in understanding how Trikafta affects cell wall composition.

Given our observations that Trikafta disrupts cell wall and cell membrane integrity, it is perhaps not surprising that we observed Trikafta-mediated phosphorylation of the *A. fumigatus* HOG pathway kinase SakA. However, two independent *sakA* null mutant strains were resistant to Trikafta-induced biomass reduction (Fig. 5A, B). Furthermore, *sakA* null mutant strains were not resistant to Trikafta effects on the inflammatory response (Fig. 6F), suggesting that the Trikafta/SakA phenotype is either downstream or independent of cell wall effects. One hypothesis we tested was that Trikafta induces an osmotic stress response in the biofilms, however, Trikafta did not increase glycerol accumulation, which is a classic response to osmotic stress (data not shown). In addition to the osmotic stress response, the HOG pathway has also been observed to be involved in general stress responses, such as those induced by oxidative stress. In *Aspergillus nidulans*, SakA has been shown to interact with other stress response proteins, such as SrkA, MpKc, and AN6892 after treatment with hydrogen peroxide (39). This general stress response reduced mitochondrial function and caused cell cycle arrest. Furthermore, the SakA homolog Hog1 mutant has been shown to have increased respiration rates and altered mitochondrial membrane potential in *Candida albicans* (40). The widely used fungicide, fludioxinil, has been shown to cause fungal cell death by hyperactivating the Hog pathway (41, 42). In the context of Trikafta treatment of biofilms, increased ion channel activation could potentially cause SakA-dependent alterations in mitochondrial function and subsequent growth arrest, which was observed in the biofilms harboring mutations in *sakA* (Fig. 5).

The fungal cell wall is critical in influencing the extent to which an immune response is generated against colonization or infection by fungal pathogens. The *A. fumigatus* cell wall contains galactomannan moieties as well as chitin and (1,3)-ý-glucan, which all play a role in triggering an inflammatory response in the host (reviewed in Beauvais and Latgé 2001). This is evidenced by early work showing that glucans can activate leukocytes, phagocytosis, and production of pro-inflammatory mediators (44). The C-type lectin receptor, Dectin-1, has been identified as the key host protein involved in recognition of fungal cell wall component (1,3)-β-glucan (45). Macrophages expressing high levels of Dectin-1 were shown to have an increased inflammatory response to *A. fumigatus*, which was completely blocked by neutralizing Dectin-1 antibodies (46). Furthermore, Dectin-1 can more efficiently recognize certain stages of *A. fumigatus* development that correspond with higher levels of β-glucans. Swollen conidia and germlings, which have the highest levels of β-glucan exposure, were shown to cause high levels of pro-inflammatory cytokine production (46). Our data demonstrate that Trikafta protects *A. fumigatus* biofilms from CFW (Fig. 6C) and causes increased susceptibility to caspofungin, which implies that there is less β-glucan content in the cell wall (Fig. 6D). This conclusion is supported by a decrease in TNFα production in the supernatants of conidia and bone marrow cells in co-culture with Trikafta compared to vehicle controls (Fig. 6E and 6F). Since this phenotype is conserved in both WT and dF508 bone marrow cells, it is likely that this is attributed to the effect of Trikafta on the fungus itself, rather than host cells. It is also possible that this effect is due to non-CFTR mediated effects (off-target) in the bone marrow cells. Further studies to determine how Trikafta affects the fungal cell wall and how this impacts host/pathogen interactions will be critical.

Our data suggest that the current CF treatment, Trikafta, causes biomass reduction, membrane permeabilization, and loss of viability in *A. fumigatus* biofilms. Furthermore, we observed that Trikafta alters *A. fumigatus* biofilm responses to cell wall stress, which correlates with reduced inflammatory cytokine production from bone marrow cells in co-culture with Trikafta treated *A. fumigatus*. To understand the implications of these observations more fully, *in vivo* models of allergic and invasive aspergillosis are necessary to define the effect of Trikafta on *A. fumigatus* biofilms in the mammalian lung.

## METHODS

### Strains and growth conditions

All strains utilized in this study are listed in Table 1. All strains were initially grown in agar (1.5%) plates containing 1% glucose minimal media (GMM; 1% glucose, 6 g/L NaNO_3_, 0.52 g/L KCl, 0.52 g/L MgSO_4_•7H_2_O, 1.52 g/L KH_2_PO_4_ monobasic, 2.2 mg/L ZnSO_4_•7H_2_O, 1.1 mg/L H_3_BO_3_, 0.5 mg/L MnCl_2_•4H_2_O, 0.5 mg/L FeSO_4_•7H_2_O, 0.16 mg/L CoCl_2_•5H_2_O, 0.16 mg/L CuSO_4_•5H_2_O, 0.11 mg/L (NH_4_)_6_Mo_7_O_24_•4H_2_O, and 5 mg/L Na_4_EDTA; pH 6.5). Conidia were collected for experiments after growth at 37 °C and 5% CO_2_ for 72h with 0.01% Tween-80 and filtered through miracloth (Millipore Sigma) to exclude hyphae.

**Table 1:**
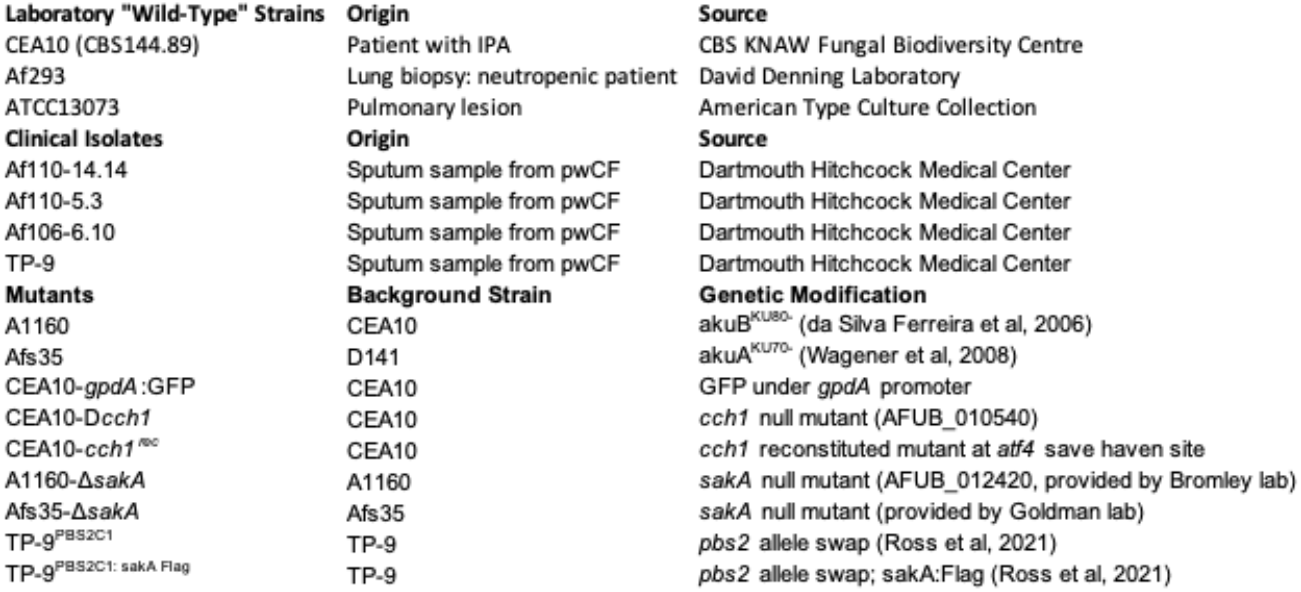
List of *A. fumigatus* strains.

**Table 2:** Trikafta treatment alters metabolic activity in select kinase KO mutants.

| AFUB # | Putative Function | % Change from vehicle | % Change from WT |
| --- | --- | --- | --- |
| AFUB_012420 | Mitogen-activated protein kinase | 176.5 | 139.9 |
| AFUB_032300 | Protein kinase, putative | 93.2 | 73.9 |
| AFUB_043130 | MAP kinase kinase Ste7 | 99.4 | 78.8 |
| AFUB_018600 | Protein kinase, putative | 105.4 | 79.9 |
| AFUB_053520 | Calcium/calmodulin dependent protein kinase, putative | 156.6 | 118.6 |
| AFUB_017750 | Protein kinase, putative | 154.8 | 117.3 |
| AFUB_096660 | Putative uncharacterized protein | 76.7 | 66.7 |
| AFUB_040390 | Protein kinase domain-containing protein | 72.6 | 63.2 |
| AFUB_020650 | Two-component osmosensing histidine kinase (Bos1) | 85.8 | 74.7 |
| AFUB_045550 | Calcium/calmodulin dependent protein kinase, putative | 86.6 | 75.4 |
| AFUB_061120 | Protein kinase domain-containing protein | 87.9 | 76.5 |
| AFUB_067080 | Putative uncharacterized protein | 90.1 | 78.4 |
| AFUB_063830 | Ubiquinone biosynthesis protein, putative | 84.0 | 73.0 |
| AFUB_101530 | Sensor histidine kinase/response regulator, putative | 91.6 | 79.7 |
| AFUB_071990 | N/A | 84.6 | 73.6 |
| AFUB_055480 | Serine/threonine protein kinase, putative | 88.2 | 76.7 |
| AFUB_098230 | Inositol kinase kinase (UvsB), putative | 91.0 | 79.1 |
| AFUB_099170 | Protein kinase Yak1, putative | 129.1 | 112.3 |

### Biofilm growth and drug treatments

For all experiments, conidia from the indicated strains were counted and seeded in liquid glucose minimal media (GMM) at 1.0X10^^5^ conidia/ml for the indicated times in 6 well plates (for biomass assays) and 96 well plates for all other assays to develop immature biofilms. Media was then removed and replaced with GMM containing either DMSO or the indicated doses of Ivacaftor (5µM=1.96µg/ml), Tezacaftor (5µM=2.60µg/ml), Elexacaftor (5µM=2.99µg/ml), or the indicated combinations (Selleckchem). In ionic stress experiments, biofilms were grown in GMM and at time of drug treatment, 500mM ionic stressors were supplemented to the media. For drug co-treatments, all biofilms were grown as above. Biofilms were treated for 1 or 3hr with the vehicle controls (dimethyl sulfoxide [DMSO] or water), voriconazole (0.25-4 µg/mL DMSO; Sigma), amphotericin B (0.125-2 µg/mL DMSO; Cayman Chemicals), caspofungin (0.015625-4 µg/mL; Cayman Chemicals), calcofluor white (6.25-50 µg/mL; Sigma), GlyH-101 (20 µM; Selleckchem), Verapamil (1mM; Sigma), SDS (0.002%; Fisher), hydrogen peroxide (0.3125-5mM; Sigma), FCCP (0.25-4µM; Sigma), Fludioxinil (31.25-500µg/ml; Sigma), antimycin A (0.25-4µg/ml; Sigma), and potassium cyanide (0.25-4mM; Sigma).

### Quantification of Submerged Biofilm Biomass

After growth in 6-well plates and treatment for the indicated times, excess supernatant was removed from the biofilms by tilting plates on side and removing pooled liquid. Biomass was then carefully scraped and collected into pre-weighed tubes. Samples were vortexed in 1ml ddH2O and then centrifuged at 15,000 rcf for 10 minutes to pellet mycelia and then repeated. Supernatants were removed, frozen, then lyophilized. Dry biomass was then quantified.

### XTT/Resazurin Assay

After biofilm treatment with drug (s) or vehicle for the indicated times, media was removed. XTT [2,3-bis-(2-methoxy-4-nitro-5-sulfophenyl)-2H-tetrazolium-5-carboxanilide] solution (0.5 mg XTT/mL 1× PBS with 25 µM menadione) (XTT sodium salt, VWR) was added at 150 µl per well and incubated at 37 °C, 5% CO_2_, and 21% O_2_ until the positive vehicle control wells were reduced (1 to 2h). Next, 100µl of the XTT solution supernatant was transferred to a 96-well plate and optical density was measured at 450 nm. For Resazurin experiments, biofilms were grown to 16hr maturity, media was removed and fresh media with DMSO or Trikafta was added. After 30 minutes, media was supplemented with 10% Resazurin and incubated for an additional five hours. Fluorescence at excitation 544 and emission 590 was used for quantification.

### Adenylate Kinase assay

To quantify alkaline phosphatase in biofilm supernatants, samples were assayed as previously described (23). Briefly, supernatants were collected and kept at 4C for no longer than 24hr. AK detection reagent (100µl: Toxilight non-destructive cytotoxicity bioassay, Lonza) was added to each well of samples (20 uL) and incubated at RT for 5 minutes prior to luminescence measurements (1hr, 5-minute intervals).

### Sytox Blue Stain of Biofilms

To assess Sytox Blue staining in treated biofilms, biofilms were grown for 16hr in filtered GMM, media removed, and treated as indicated. After treatment, biofilms were stained with Sytox Blue (Thermo Scientific) 1:1000 for five minutes and then imaged using a Nikon spinning disc confocal microscope at 20X magnification. Images were processed and analyzed using the Fiji image analysis software (47). Raw image stacks were resliced from top down without interpolation and maximum projected. The Fiji plugin JaCoP was used to analyze the co-localization of the cytosolic GFP signal with the Sytox signal (48). In the JaCoP plugin the Sytox channel was set to channel 1 and GFP was set to channel 2. Channel thresholds were set manually to exclude the background signal. The Manders’ overlap was quantified between the two channels and M2 was recorded as this represents the fraction of GFP positive pixels that overlapped with SYTOX positive pixels as a read out for level of Sytox staining across conditions (49).

### Phospho-sakA Western Blot

To assess phosphorylation of SakA by Western Blot, a protocol for submerged biofilms was adapted from stationary cultures (26). Briefly, submerged biofilms (TP9^PBS2C1:sakA-Flag^) were grown in 6 well plates for 16h, media removed, and indicated treatments in GMM were added for an additional 30 minutes. Biomass was scraped into 2ml screw cap tubes, frozen, and lyophilized. They were then bead beaten in 1ml protein extraction buffer and total protein was quantified by Bradford kit. For Western, 40μg protein was used. Total protein was transferred from a 10% SDS-PAGE gel onto a nitrocellulose membrane using the Trans-Blot turbo transfer system (Bio-Rad). Total Flag-sakA was detected using the Flag M2 antibody 1:2000 (Sigma) and phosphorylated SakA was detected using anti-P-p38 antibody 1:1000 (Cell Signaling). To quantify fluorescence, imaging was performed using the LI-COR Odyssey CLX System according to manufacturer’s protocols Total Flag and phospho-sakA were normalized to total protein using the REVERT total protein stain (LI-COR Biosciences).

### Bone Marrow Co-Culture

Bone marrow cells were collected from the tibia and femur of female C57Bl/6 WT and dF508 mice (10-12 weeks). The dF508 and littermate control breeders were obtained from Case Western Reserve University (Cystic Fibrosis Mouse Models Core). Bones were removed and BM cells isolated by flushing bone marrow out using DMEM-based tissue culture media. Cells were pelleted, red blood cells were lysed, and cells were set up in co-culture with the indicated strains of conidia at an MOI of 1:10 supplemented with 20ng/ml IFNγ for 16hr. Media was collected and assayed for total TNFα concentration according to manufacturer’s instructions (R&D systems).

### Statistics

All experiments were performed with at least three biological replicates and data points are averages of technical replicates for each individual experiment. For experiments comparing two groups, a standard student’s t test was performed. For experiments comparing two groups under multiple conditions, a 2-way ANOVA with Sidak’s multiple comparisons was used or Dunnett’s multiple comparisons was used when comparing to one control. When comparing more than two groups a one-way ANOVA with Tukey’s multiple comparisons was used. All statistical analyses were performed using GraphPad Prism 9.4.1.

## STUDY APPROVAL

Murine studies for collection of bone marrow cells were approved by the Dartmouth Institutional Animal Care and Use Committee (IACUC) under protocol number 00002167.

## DATA AVAILABILITY

All underlying data are available in either supplement or at author’s request.

## AUTHOR CONTRIBUTIONS

J.T.J. designed experiments, performed the majority of experiments, performed data analysis, generated figures, and wrote the manuscript. K.A.M. designed Sytox Blue and ion channel inhibitor experiments and performed microscopy. E.M.V. performed AK experiments, data analysis, and figure layout design. C.T.S.P. performed analysis on Sytox Blue/GFP images, contributed to figure layout, and contributed to writing the manuscript. C.K.P. generated and confirmed CEA10-Δ*cch1* and CEA10-*cch1^rec^* strains. B.S.R. isolated and identified *A. fumigatus* clinical isolates and performed Western Blot analysis. N.R and M.J.B. generated the kinase knockout library collection. RAC conceived/funded the study, designed experiments, and wrote the manuscript.

## ACKNOWLEDGEMENTS

This work was supported by the efforts of R.A.C. through funding by a Cystic Fibrosis Foundation research award (CRAMERGO19). KAM was supported by NIH award T32-AI007519. Core facility support provided by NIH grant P20-GM113132 to the Dartmouth BioMT COBRE. Additional support was provided by the Cystic Fibrosis Foundation Research Development Program (STANTO19R0) and NIH P30-DK117469 (Dartmouth Cystic Fibrosis Research Center).

